# Inhibiting presynaptic calcium channel mobility in the auditory cortex suppresses synchronized input processing

**DOI:** 10.1101/2022.03.30.486338

**Authors:** Katrina E. Deane, Ruslan Klymentiev, Jennifer Heck, Melanie D. Mark, Frank W. Ohl, Martin Heine, Max F. K. Happel

## Abstract

The emergent coherent population activity from thousands of stochastic neurons in the brain is believed to constitute a key neuronal mechanism for salient processing of external stimuli and its link to internal states like attention and perception. In the sensory cortex, functional cell assemblies are formed by recurrent excitation and inhibitory influences. The stochastic dynamics of each cell involved is largely orchestrated by presynaptic CAV2.1 voltage-gated calcium channels (VGCCs). Cav2.1 VGCCs initiate the release of neurotransmitters from the presynaptic compartment and are therefore able to add variability into synaptic transmission which can be partly explained by their mobile organization around docked vesicles. To investigate the relevance of Cav2.1 channel surface mobility for the input processing in the primary auditory cortex (A1) *in vivo*, we make use of a new optogenetic system which allows us to acutely cross-link Cav2.1 VGCCs via a photo-cross-linkable cryptochrome mutant, CRY2olig. In order to map neuronal activity across all cortical layers of the A1, we performed laminar current-source density (CSD) recordings with varying auditory stimulus sets in transgenic mice with a citrine tag on the N-terminus of the VGCCs. Clustering VGCCs suppresses overall sensory-evoked population activity, particularly when stimuli lead to a highly synchronized distribution of synaptic inputs. Our findings reveal the importance of membrane dynamics of presynaptic calcium channels for sensory encoding by dynamically adjusting network activity across a wide range of synaptic input strength.

**Statement of Significance:** Voltage Gated Calcium Channel (VGCC) mobility plays an important role in neuronal firing dynamics. Failure of these channels to function or be regulated has been linked to migraine and ataxia. We here link the microscopic process of VGCC mobility to the mesoscopic population dynamics as a mechanism to regulate and appropriately amplify synaptic inputs of different strengths to the mouse primary auditory cortex. We also demonstrate a novel and effective technique with which VGCC function can be further explored in meso- or macroscopic scales and with behaving subjects. We believe that this is of importance to the broader scientific community in aspects of non-linear scaling in the brain, potential translational applications, and basic research on cortical mechanisms of physiological function.

## Introduction

The primary auditory cortex (A1) receives par-processed spectral inputs and subsequently categorizes sound and initiates auditory-guided behaviors (King et al. 2018; Nelken 2020; Ohl 2015). Coherent population activity within the A1 is primarily generated via thalamocortical input and intracortical recurrent microcircuits (Happel et al. 2010; Liu et al. 2007) and recruited by sensory inputs (Beltramo et al. 2013). Excitatory and inhibitory postsynaptic dynamics are considered to be a major origin for such input-derived population activity in the sensory cortex (Fu et al. 2014; Gabernet et al. 2005; Wu et al. 2008). Cav2.1 channels, an important subtype of voltage-gated calcium channels (VGCC) triggering action potential-mediated synaptic vesicle release, are important regulators of neuronal dynamics and communication (Heck et al. 2021).

It was demonstrated *in vitro* that synapse-specific release properties depend on the molecular lateral mobility of Cav2.1 channels within the presynaptic membrane (Heck et al. 2019) and tightly time calcium-mediated synaptic vesicle release can encode sensory information (Hay and Segev 2015; Young and Veeraraghavan 2021). It has therefore been suggested that the regulation of presynaptic VGCCs mobility and vesicle release-sites would control synaptic release probability and short-term plasticity (Böhme et al. 2018; Heine et al. 2020).

Neurons across A1 cortical layers exhibit several sensory-evoked activities, reflecting the flow of information through cortical circuits (Harris and Mrsic-Flogel 2013; Sakata and Harris 2009). Afferent inputs mainly recruit recurrent microcircuits in granular layers (Hackett et al. 2011) and thereby yield highly synchronized synaptic inputs. Supragranular layers, which densely connect across the neocortex, and infragranular layers, which receive secondary thalamic input, mediate corticocortical connections in the service of, for instance, spectral integration, corticocortical integration, temporal processing, and corticothalamic feedback (Francis et al. 2018; Happel et al. 2014; Jeschke et al. 2021; Moeller et al. 2010).

Exactly how membrane motility of presynaptic VGCCs may influence the gating of afferent inputs in the sensory cortex at a population level is yet elusive. Therefore, we targeted the N-termini of Cav2.1 channels in the right A1 of transgenic knock-in mice, Cacna1a^Citrine^ (Mark et al. 2011), with an optogenetically aggregating cryptochrome mutant, CRY2olig (Heck et al. 2019; Taslimi et al. 2014), via a feed-back-controlled anti-GFP intrabody. We recorded local field potentials *in vivo* across A1 cortical layers under ketamine anesthesia and computed current source density (CSD) profiles (Brunk et al. 2019; Deane et al. 2020; Happel et al. 2010) before and after optogenetically-induced VGCC clustering.

We compared responses to two different kinds of auditory stimulus sets that reflect different aspects of spectral and temporal auditory processing: click trains and amplitude modulated (AM) tones. Clicks are characterized by a broad energy spectrum covering the hearing range of mice and hence activate hair cells along the entire basilar membrane (Lu and Wang 2000). Click trains thereby cause repetitive and highly synchronized afferent thalamocortical synaptic inputs over a broad tonotopic area of the A1. On the other hand, AM tones have a narrow energy spectrum, the amplitude of which periodically varies in time. Contrasting responses to these two types of spectral energy should reveal key differences in how a population with clustered VGCCs would internally synchronize and respond to certain specific aspects of sounds.

We found that light-induced aggregation of Ca_V_2.1 channels in A1 generally suppressed sensory-evoked synaptic population activity across all cortical layers. Particularly, click stimuli that lead to a highly synchronized distribution of synaptic inputs in thalamocortical input layers IV and V, showed a significant reduction. Effects on less synchronized input-derived AM-evoked responses were more subtle. In control groups we found the reversed effect, which may be explained by heat from the superficial laser illumination (Arias-Gil et al. 2016). Our study reveals the importance of the membrane motility of VGCCs to support the gain function of cortical recurrent excitation. Presynaptic membrane dynamics thereby facilitate population activity across a wide range of incoming synaptic input, which may be a critical network characteristic for adaptive and ongoing sensory encoding.

## Materials & Methods

### Ethical approval and subjects

Experiments were conducted in accordance with ethical animal research standards defined by German Law and approved by an ethics committee of the State of Saxony-Anhalt under license 42502-2-1394LIN. All experiments were carried out with adult male mice (*Mus musculus*, 8-13 weeks of age, 18-28 g body weight, total *n* = 27) of the transgenic line C57BL/6J *Cacna1a*^*Citrine*^ (Mark et al. 2011). Note that female animals were not used as possible variances due to sex was not in the scope of our study.

### Optogenetic cross-linking of Cav2.1 calcium channels in vivo

The knock-in mouse line used in this study expresses a Citrine tag, a YFP/GFP derivate, at the N-terminus of Cav2.1 voltage-gated calcium channels, which has been reported to be specifically detected by GFP antibodies (Mark et al. 2011). Here, we used a recently developed system that utilizes a feedback-controlled intracellularly expressed anti-GFP nanobody to target the Citrine tag and at the same time equip the Cav2.1 N-terminus with a photo-cross-linkable cryptochrome mutant, CRY2olig (Taslimi, Justin D Vrana, et al. 2014). Under blue (477-488 nm) light exposure, CRY2olig reversibly snaps together (Schematized in Figure 1F). As previously shown, CRY2olig reaches ∼60 % clustering immediately after light stimulation and clusters decrease to ∼30 % over 30-40 minutes and to ∼0 % again in the duration of 160 minutes (see Heck et al. 2019; Taslimi et al. 2014)

**Figure 1.**
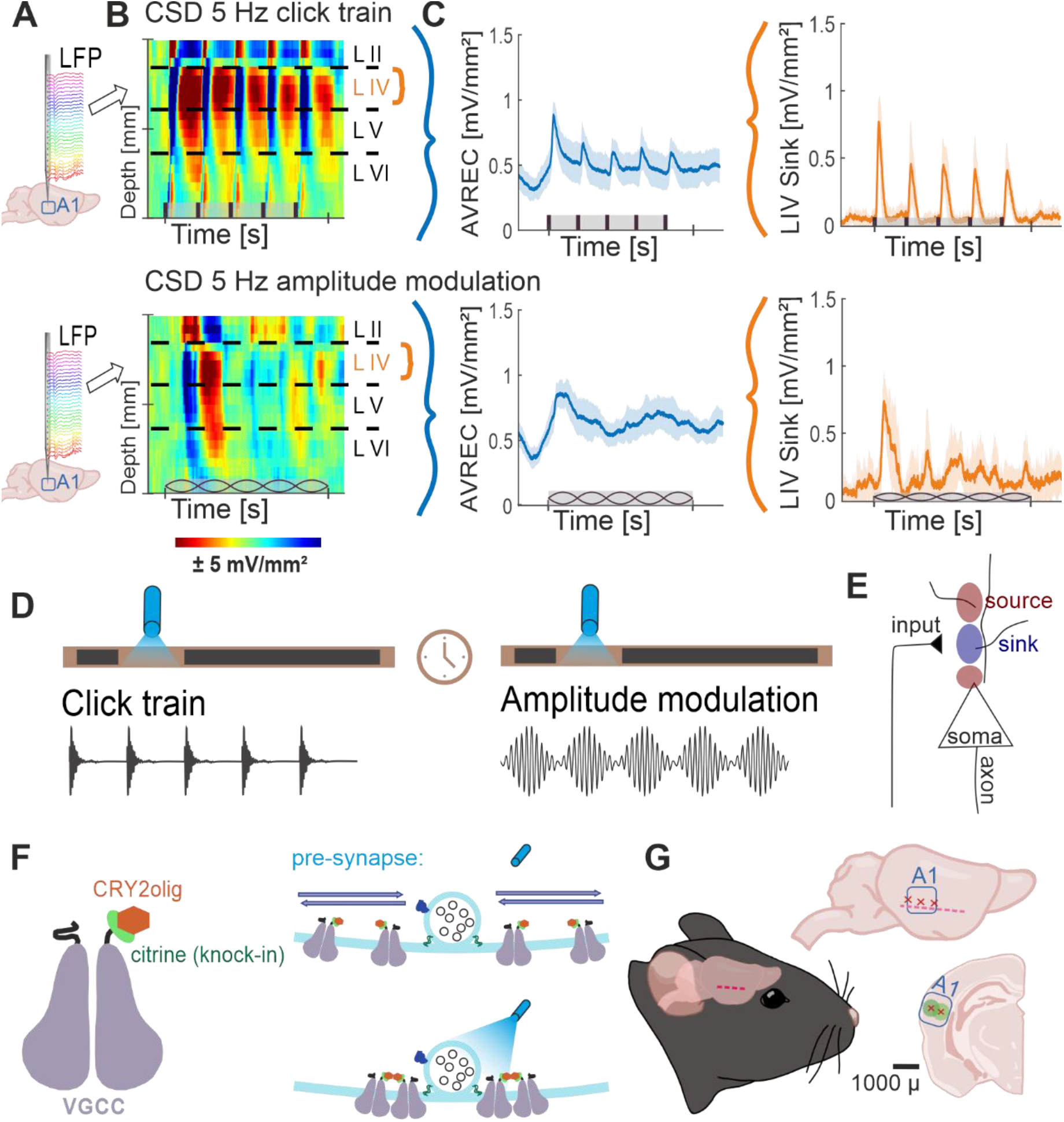
Project Flow; **A**. Representation of 32 channel shaft electrode implanted perpendicularly into the auditory cortex, recording local field potentials (LFPs) over time. **B**. CSD profiles of cortical response to 5 Hz click train (top) and amplitude modulation (bottom) click trains. CSDs show neuronal activity down the cortical depth of the auditory column over time. **C**. Average rectified (AVREC) CSD traces (left) and layer IV sink traces (right) for CSD profiles in B. **D**. Protocol of measurements Click Trains and Amplitude Modulation before and after blue light laser stimulation. **E**. Schematic of movement of ions recorded by current source density (CSD) analysis. Input to the neurons causes intake of positive ions, creating negatively charged extracellular space: sink. Sources are the balancing loop current created by depolarization of the cell and the sink. **F**. Schematic representation of a voltage-gated calcium channel (VGCC) with a knock-in citrine tag on its N-terminus and a CRY2olig protein attached (left) and of the normal movement of VGCCs in the pre-synapse around a docked vesicle with no laser light (top right) compared to the clustering of several VGCCs around the docked vesicle with the laser light on (bottom). **G**. Schematic representation of lentivirus transduction: cut made between ear and eye, muscle scraped gently down, three trepanations and subsequent transductions across the A1, ∼1 mm apart, at two depths, 300 and 600 μ, where, at each, 23 nl was transduced 9 times every 3 seconds.

### Anesthesia

Ketamine–xylazine was administered at surgery onset and throughout the acute experiment to maintain a steady level of anesthesia. Infusion of 20% v/v ketamine (ketavet or ketabel) (50 mg ml−1, Ratiopharm GmbH, Ulm, Germany), 5% v/v xylazine (Rompun 2%, Bayer Vital GmbH, Leverkusen, Germany), and 75% v/v of isotonic sodium chloride solution (154 mmol 1−1, B. Braun AG, Melsungen, Germany) was given intraperitoneally for an initial dose of 4 ml per 1 kg of body weight. A needle was placed subcutaneously or intraperitoneally to maintain anesthetic status with an infusion rate of ∼0.4 ml per 1 kg of bodyweight per 1 hr during the experiment. Anesthetic status was regularly checked (every 7.5–10 min) by paw withdrawal reflex, tail pinch, and breathing frequency. Body temperature was kept stable at 37°C.

### Transduction of virus

Surgical transduction of the lentivirus in the A1 was performed as follows. Ketamine-xylazine was administered once intraperitoneally as described above and the temporal bone was exposed via a single 5mm cut from in front of right ear to behind the right eye and gentle removal of the temporal chewing muscle by scraping it down off the bone for 2 mm. Three ∼0.5 mm holes were created above the A1 1 mm below the temporal suture through trepanation, 1 mm apart (Figure 1G). Virus, containing Cry2olig, was injected 300 and 600 μm deep at each of the three sites. Each of the 6 injection sites received 23 nl of virus 9 times every 3 seconds, totaling 207 nl of virus at each site and 1,242 nl across the A1. This virus, an LV-CAG which contains the sequence: CIBN-Xm233-EOS-CCR5, did not contain a fluorescent protein and could not be stained in immunohistochemistry to show the spread (see Rieder et al. 2015, Figure 5). Lentiviruses have been shown as efficient in their role of infecting neurons *in vivo* in, amongst other species, rats (Naldini et al. 1996a; Naldini, et al. 1996b). While the spread of the virus is limited, it has been shown that 200 nl injection volumes diffuse within a spherical region with a diameter around 200-600 μm (Desmaris et al. 2001; Osten et al. 2006 see Figure 13.3). Therefore, between each subject, we can assume a large coverage of the A1 down the depth of the cortical column. The vehicle control group underwent the same procedure and injected with an equal volume of a lentivirus not containing Cry2olig. The naïve control group received no surgery prior to electrode implantation (see below). After surgical injection, the muscle was gently placed over the trepanated temporal bone, and the skin was sutured. Metacam (2 mg/ml, Boehringer Ingelheim, Ingelheim am Rhein, Germany) was administered at the onset of surgery subcutaneously for a dose of 1 ml per 1 kg of bodyweight and two consecutive days peri-operation for a dose of 0.5 ml per 1 kg of bodyweight.

The virus was allowed to express for 4 weeks before auditory recording. The implantation and recording were carried out on the same day, per subject, so it was necessary to perform two surgeries to allow time for expression.

### Electrode placement and recording

The surgical procedure for electrophysiological recording has been previously described in detail in Mongolian gerbils (Deliano et al. 2020). With the same surgery for mice, briefly, ketamine–xylazine was administered as described above and the right auditory cortex was exposed by removal of the temporal chewing muscle and trepanation of a 3×4 mm opening in the temporal bone between the ear and eye. A small hole was drilled on the contralateral hemisphere for implanting a stainless-steel reference wire (Ø 200 μm). Animals were head-fixed in a Faraday-shielded acoustic soundproof chamber with an aluminum bar, affixed by UV-curing glue (Plurabond ONE-SE and Plurafill flow, Pluradent, Offenbach, Germany). The dura was cut and a 32 channel tungsten electrode (A1×32-50-413, NeuroNexus, Ann Arbor, MI, USA) was implanted perpendicularly into the right A1, located via vascular landmarks (Brunk et al. 2019; Happel et al. 2010). Animals were sacrificed by decapitation at the end of the 6–8-hour experiment.

Note that the surgery took place in white light necessarily which may have caused some VGCC clustering in CRY2olig-treated animals. The setup was in total darkness except for periods of red light after electrode implantation and an hour of baseline recording was taken in part to allow clustering to relax before taking pre-laser measurements (described below).

### Auditory stimuli and blue light laser

A speaker was located 1 m posteriorly (Tannoy arena satellite KI-8710-32, Tannoy, London, UK) to the head-fixed mice. Stimuli were generated in Matlab (R2006b, The Mathworks, Natick, MA, USA), converted into analog (sampling frequency 1000 Hz, NI PCI-BNC2110, National Instruments, Austin, TX, USA), routed through an attenuator (g-PAH, Guger Technologies, Graz, Austria), and amplified (Thomas Tech Amp75, Tom-technology, Ilirska Bistrica, Ljubljana). A microphone and conditioning amplifier were used to calibrate acoustic stimuli (G.R.A.S. 26AM and B&K Nexus 2690-A, Brüel & Kjær, Naerum, Denmark).

Three types of stimuli were provided during recording. The first was tonotopy: a series of pseudo-randomized pure-tone frequencies covering a range of seven octaves with considerable sound pressure levels (cf. Happel and Ohl 2017; tone duration: 200 ms; tone frequency: 125 Hz to 32 kHz; inter-stimulus interval: 800 ms; 50 pseudorandomized repetitions; 65 dB sound pressure level; 7.5 min per measurement). We determined the best frequency (BF) as the frequency evoking the strongest response in the averaged granular CSD channels (see below; see Supplementary Figure 3 Zempeltzi et al. 2020). The second was click train measurement (Figure 1D): a series of pseudo-randomized presentation-frequency noise-click trains with a carrier frequency of the determined BF (stimuli duration: 999 ms; click presentation-frequency: 5 and 10 Hz; inter-stimulus interval: 500, 200, 100, 50, and 25 ms respectively; carrier tone: BF; 30 pseudorandomized repetitions before the laser and 50 after; 90 dB sound pressure level; 10 min before the laser and 15 min after per measurement). The third was an amplitude modulation measurement (Figure 1D): a series of pseudo-randomized frequency modulations of a tone at the determined BF (stimuli duration: 999 ms; modulation frequency: 5 and 10 Hz; carrier tone: BF; inter-modulation interval: 500, 200, 100, 50, and 25 ms respectively; 30 pseudorandomized repetitions before the laser and 50 after, 65 dB sound pressure level, 10 min before the laser and 15 min after per measurement).

At the onset of recording, a light fiber was situated 5 mm above the cortical surface and directed to shine light over the 3×4 mm opening over the A1. The fiber output 5 mW of power at 477 nm wavelength and was switched on for 20 seconds after all pre-laser measurements were taken. Light intensity propagated through the entire cortical depth and, henceforth, led to the aggregation of Cav2.1 channels within the A1.

The protocol was as follows: surgery (described above), baseline tonotopy recording for 1 hour, one 10-minute amplitude modulation measurement, 30 minutes of tonotopy, one 10-minute click train measurement, 30 minutes of tonotopy, blue light laser presentation (5mW; 20 s; 477 nm), 1 hour of click train measurements, one tonotopy, blue light laser, 1 hour of amplitude modulation measurements, and one tonotopy. Spontaneous activity was recorded for 2 minutes before and after each laser presentation.

### Electrophysiological recording

Recorded LFPs (Figure 1A) taken during the above stimuli presentation from the NeuroNexus electrode were fed via an Omnetics connector (HST/32V-G2O LN 5V, 20× gain, Plexon Inc., Dallas, TX, USA) into a PBX2 preamplifier (Plexon Inc.) to be pre-amplified 500-fold and band-pass filtered (0.7– 300 Hz). Data were then digitized at a sampling frequency of 1000 Hz with the Multichannel Acquisition Processor (Plexon Inc.).

### Current source density profiles and average rectification

Based on the recorded laminar local field potentials, the second spatial derivative was calculated using Matlab (R2016a), yielding the current source density (CSD) distribution (Figure 1B, Schematized in Figure 1E) as given by eqn (1):

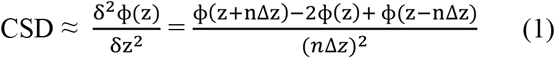

Where ϕ is the electric local field potential, *z* is the spatial coordinate perpendicular to the cortical laminae, Δ is the sampling interval, and *n* is the differential grid (Mitzdorf 1985). LFP profiles were smoothed with a weighted average via Hamming window of seven channels, corresponding to a spatial kernel filter of 300 μm (Happel et al. 2010). Current sinks in the CSD distribution correspond to the activity of excitatory synaptic populations due to the local spatiotemporal current flow of positive ions from extracellular to intracellular space, while current sources mainly reflect balancing return currents evoked by this extracellular hyperpolarization.

CSD profiles were further transformed by averaging the rectified waveforms of each channel (Figure 1C, left) as seen in eqn (2):

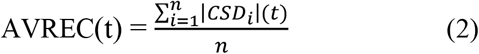

where *n* is the number of recording channels and *t* is time. This average rectified CSD (AVREC) is a measure of the overall local current flow of the columnar activity (Givre et al. 1994; Schroeder et al. 1998).

Based on tone-evoked CSD distributions, we assigned the main sink components to the cortical anatomy as follows: the early dominant sink components are attributed to lemniscal thalamocortical input, which terminates in cortical layers IV and the border of V and VI. Note that in our previous work, CSDs have been shown from Mongolian gerbils which have a thicker A1 and subsequently more distinguishable layer sink components (Deane et al. 2020; Happel et al. 2010); in the mouse A1, we will be distinguishing layer I/II (as II), III/IV (as IV), V, and VI as primary sink component layers based on mouse A1 CSD from (Yamamura, Sano, and Tateno 2017). Each layer was transformed into layer sink traces (Figure 1C, right; cf. Zempeltzi et al. 2020) by averaging sink activity of each channel attributed to the layer as seen in eqn (3):

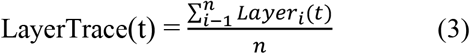

where *Layer* } x<0, *n* is the number of individual channels attributed to the layer, and *t* is time in milliseconds. This layer trace gives us the temporal local current flow of sink activity per cortical layer for which it is calculated.

### Statistical analysis

After calculation of CSDs and traces, the root mean square (RMS), a metric of strength of activity, was calculated along each trial trace within the time window: stimulus onset to 1000/Hz ms (200 ms for 5 Hz stim). Peaks were also detected during this time window; highest peak prominence was selected as the relevant peak feature selected for that trial after it crossed an arbitrary threshold of peak prominence of at least 0.00008 (according to the findpeaks function in Matlab). This allowed us to exclude trials where no cortical activity after the stimulus onset was detected. Peak latency and amplitude were recorded along the first time window from stimulus onset and in each following window of the same duration across the stimulus (0-200, 20-400, …, 800-1000 ms for 5 Hz). RMS and peak amplitude features are comparable as metrics of cortical strength and therefore much of the analysis was computed with RMS unless peak latency was also relevant. For spontaneous measurements, RMS was calculated in 1400 ms time bins and no peaks were detected.

Data was analyzed using linear mixed models (LMMs) due to the presence of repeated measurements within subjects which could be dealt with using random effects structure. LMMs have several advantages, such as dealing with missing values and ability to add various configurations of random effects, such as crossed or nested (Alday, Schlesewsky, and Bornkessel-Schlesewsky 2017). Full description of LMMs is beyond the scope of this paper, but readers can refer to the papers (Alday et al. 2017; Harrison et al. 2018).

LMMs were implemented using R (version 3.6.1) and the *nlme* package (Pinheiro et al. 2021). The dependent variable, *RMS*, was log-transformed in order to meet the model assumptions (normality of residuals in particular). Independent categorical variables, *Group* (three levels: naive control, viral control and CRY2olig-treated) and *Measurement* (two levels: pre-laser and post-laser) were encoded using treatment coding and were added in the model with an interaction. CRY2olig group and post-laser Measurement were selected as an intercept and each level of the categorical variables was contrasted to it (for example, Control group and pre-laser *Measurement* versus CRY2olig group and post-laser *Measurement*). Separate models for each combination of signal frequency (5 Hz, 10 Hz) and layer (AVREC, I_II, IV, V, VI) were built in a following structure:

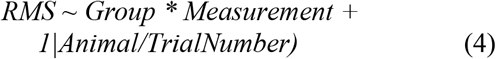

where *Group* and *Measurement* are the fixed effects and *Animal* and *TrialNumber* are the random effects. Random effect for *Animal* controls for the assumption that each animal has a different baseline activity, while the rate of change (slope) is the same. Nested random effect (*Animal/TrialNumber*) controls for the dependence of trials for the same animal and assumption that observations between trials for the same animal are more similar to one another than to trials from the other animals. In this way, variability of the same animal carries less weight on the outcome.

Bonferroni corrected (n=14) Student’s *t* tests were further calculated to show specific between-group differences both before and after the laser for each layer trace and AVREC RMS (Table 3) at a raw, single-trial level. Bonferroni corrected Student’s *t* tests were also performed within-group from pre- to post-laser on all traces’ RMS values (Table 4). Corresponding to each Student’s *t* test, Cohen’s d effect size was computed to gauge the magnitude of difference between the groups, or the strength of relationship of the dependent to independent variables. A Cohen’s d effect size of medium, for example, would mean that ∼70 % of the control group fell below the experimental group mean. *p* value results from single-trial *t* tests are best interpreted in conjunction with effect size results.

To test for synchronicity, we computed vector strength. This was done by matching the latency of the most prominent peak in each time window across the duration of the stimulus to the phase of the amplitude modulation at that those time points. Each phase result was considered a unit vector with the orientation of the given phase. Vector strength was the resultant length of summing those unit vectors (c.f. Middlebrooks, 2008). This analysis was performed on the AVREC of each click train and amplitude modulation of 5 and 10 Hz. Note that click trains do not have a phase as they are not an extended, modulating tone. Therefore, synchronicity of the click train results was calculated using the supposed phase of an AM tone at those latencies. Four 15-minute measurement were recorded after the laser presentation and vector strength was calculated for each of them. With this, we compared the vector strength of groups and measurements as factors in an ANOVA.

## Results

### Current source density profiles and their AVREC and layer traces show a reduction of cortical activity after clustering in the CRY2olig group

To qualitatively understand cortical response in the A1, Figure 2 displays the grand-averaged CSD profiles of the CRY2olig group and the naïve control group before and after laser presentation in response to click trains and amplitude modulation, each at 5 Hz. The click train stimulus has an immediate and repetitive burst of broad spectral energy across all frequencies of the energy spectrum, while the spectral energy of AM tones is fluctuating in amplitude by a modulation frequency but constant at a single carrier frequency consisting of a narrow energy spectrum. The bursts of energy from repetitive clicks, which quickly recruit all of the full tonotopic map of the auditory pathway (Liu et al. 2019; Lu and Wang 2000), create impulse following responses: a strong cortical recruitment of population response to each consecutive stimulus. A single impulse response—a strong and timely cortical recruitment—can be seen after the onset of the AM tones after a delay due to the ramping-up time of 100 ms from 5 Hz modulation.

**Figure 2.**
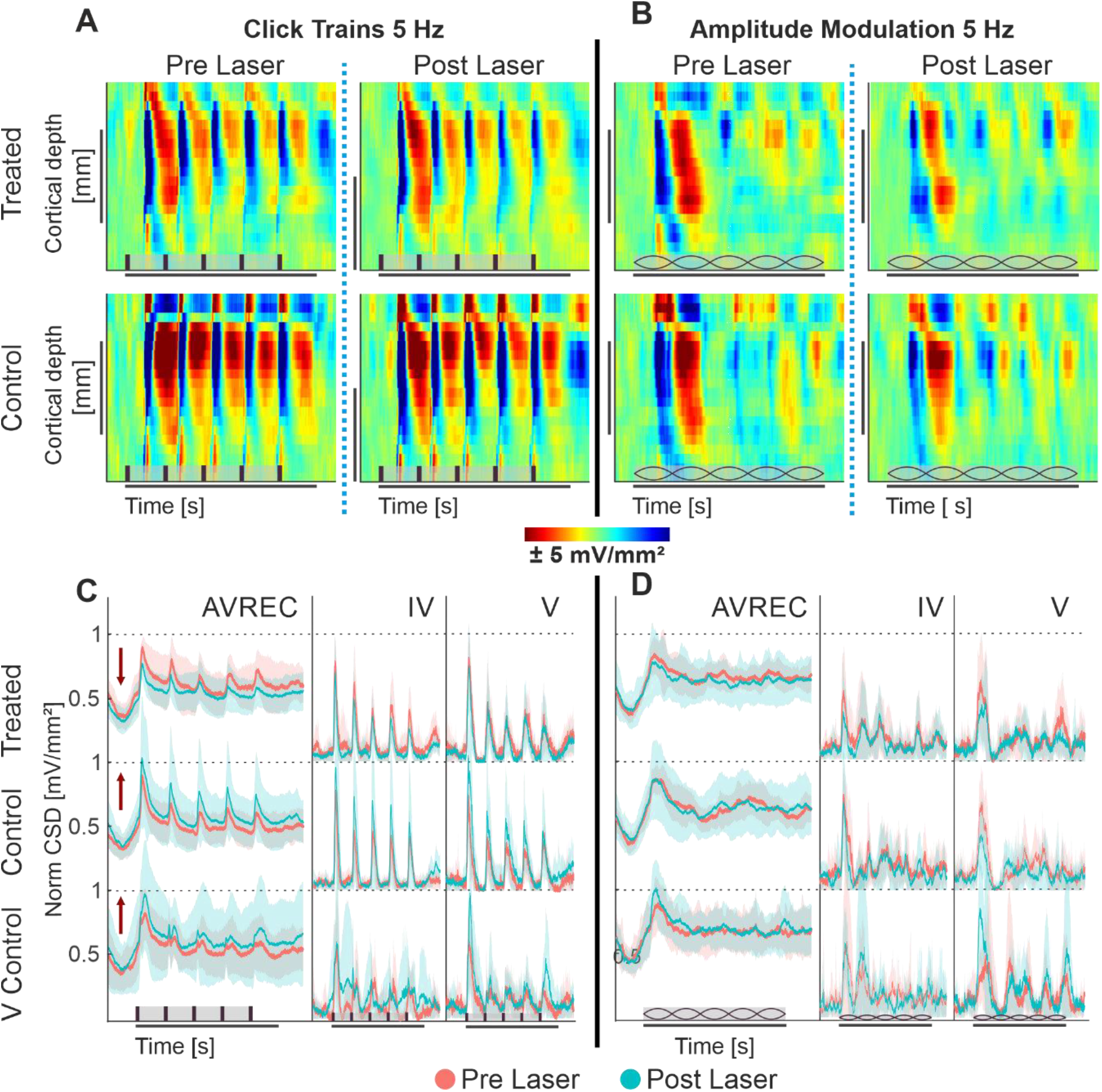
A-B: Grand averaged current source density profiles of CRY2olig-treated and naïve control group; **A**. CSD profiles of the auditory cortical column in response to 5 Hz click train in the CRY2olig group (top row) and naïve control group (bottom row) before laser presentation (left column) and after (right column). **B**. CSD profiles of the auditory cortical column in response to 5 Hz amplitude modulation of CRY2olig and naïve control groups before laser presentation and after. CSD is set in space—depth—over time with color representing the strength of activity. Blue areas are sink and represent population activity. Stimulus type represented in brown on the x axes: vertical bars for clicks and lemons for amplitude modulation. **C-D: AVREC and layer traces (±STD);** Some layer traces not included. **C**. 5 Hz click train AVREC, layer IV, and layer V (left to right column) traces for CRY2olig-treated, naïve control, and viral control groups (top to bottom rows). Pre-laser measurements are peach, post-laser measurements are light blue. SEM is shown in semi-transparent peach or blue, respectively. **D**. 5 Hz amplitude modulation AVREC, layer IV, and layer V traces for CRY2olig and control groups. Pre- and post-laser measurements are in peach and blue, SEM shown in semi-transparent peach and blue respectively. Stimuli type is represented in brown on the x axes: vertical bars for clicks and lemons for amplitude modulation.

Under ketamine anesthesia, we found that population activity within the auditory cortex of the mouse was able to clearly follow 5 Hz click trains with corresponding impulse responses (Figure 2A). Following the laser, there was a noticeable decrease in cortical activity per impulse response in the CRY2olig group but not in controls. This is also very clearly shown in Figure 2C, as explained below.

Amplitude modulation at 5 Hz (Figure 2B), did not drive an auditory following response nor as strong of an initial impulse response. Rather the cortical response represented a less categorically clear current source density profile in response to the constant fluctuating spectral energy of the stimulus.

CSD profiles were transformed into AVREC traces, or the overall columnar response, and layer traces, sink activity from individual layers, (see Figure 1) to show the overall and layer activity in the cortical column, respectively. Consistant with the findings from CSD profiles (see Figure 2A-B) we found that 5 Hz click trains evoke a clear impulse following response in the AVREC and layer traces (Figure 2C). This is true for all three subject groups. The controls differ from the CRY2olig group here, however, in directionality from pre- (peach) to post- (blue) laser measurements. While there is a general increase found in cortical activity and in the layers after the laser presentation in control groups, an overall suppression of activity after laser presentation dominates the CRY2olig group.

The cortical response to 5 Hz amplitude modulation is shown in Figure 2D. The initial cortical response to the onset of the modulated tone does not rise to a sharp peak, such as in response to clicks, due to the slow ramping of amplitude (100 ms to peak amplitude) and there is no following response in the AVREC or the layers. For the amplitude modulation, there are no clear differences pre- and post-laser across the CRY2olig and the naïve control groups but the viral control group does display a similar increase in activity after the laser.

### Strong cortical recruitment and exaggerated recurrent excitation is more susceptible to influence over population dynamics

#### Linear mixed models show that the suppression after the laser is consistent across both click trains and amplitude modulated tones for the CRY2olig group

We calculated LMM as a conservative measure to detect overall effects, due to the presence of repeated measurements within subjects. The intercept used was the CRY2olig group post-laser, meaning that comparisons run were: CRY2olig group pre-laser vs post-laser and both control groups post-laser vs the CRY2olig group post-laser (Table 1). Figure 3A and B show the effect plot results of the LMM for click trains of 5 Hz (3A) and amplitude modulation of 5 Hz (3B) across all layers for the 3 groups pre- and post-laser. There was a highly significant difference between the CRY2olig group pre- and post-laser in the AVREC, layer IV, and V of the click trains with a downward trend, indicating a significant decrease in activity after laser-induced clustering. The only exception is a significant increase in activity after the laser in layer VI. Suppression after the laser was also highly significant in the AVREC of the amplitude modulation comparison and there was also significant suppression in layers II, IV, and VI in amplitude modulation measurements.

**Figure 3.**
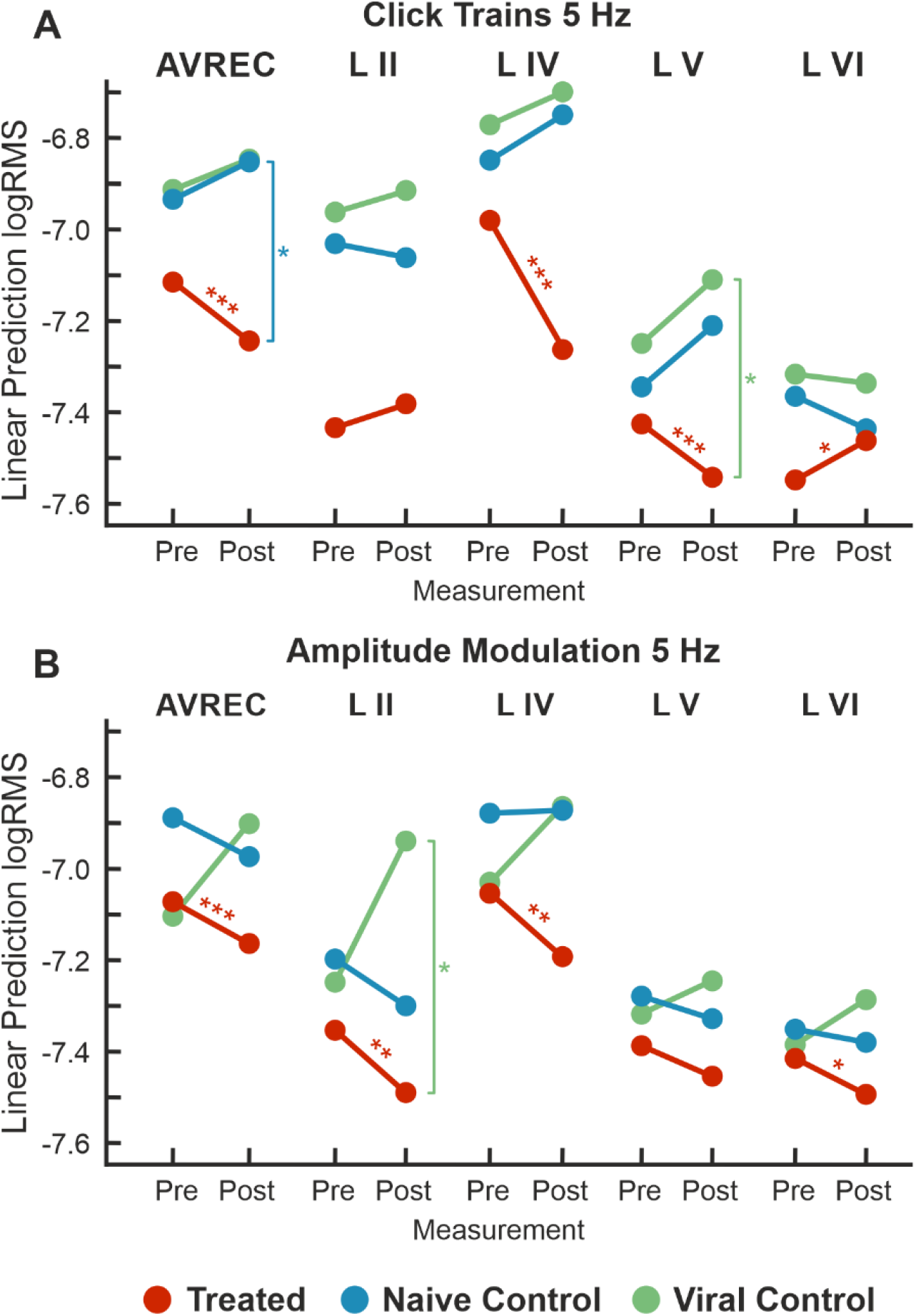
LMM effects for 5 Hz; **A**. LMM effect plots for linear prediction over measurement (pre- to post-laser) for CRY2olig-treated (orange), naïve control (blue), and viral control (green) groups across the AVREC and all cortical layers in response to click trains. **B**. LMM effect plots for linear prediction over measurement (pre- to post-laser) for CRY2olig, naïve, and viral control groups across the AVREC and all cortical layers in response to amplitude modulation. Results for LMM comparisons—CRY2olig pre-vs treated post-laser (orange), CRY2olig vs naïve post-laser (blue), and CRY2olig vs viral post-laser (green)—are overlaid as significance stars. p < 0.05 *, <0.01 **, <0.001 ***.

**Table 1.** LMM results for Click train stimulus: Comparisons run with the logRMS and the Intercept: CRY2olig-Treated:Post-Laser. Therefore the intercept, treated vs Naïve control post-laser, treated vs viral control post-laser, and treated pre-laser vs post laser are compared in the AVREC and layer traces. Significant results, p<0.05, are in bold.

| Predictors | Layer | Estimates | SE | Statistics | df | p |
| --- | --- | --- | --- | --- | --- | --- |
| Treated vs Naïve-Control : Post-Laser | **AVREC** | 0.397 | 0.178 | 2.235 | 24 | **0.035** |
|  | II | 0.327 | 0.222 | 1.475 | 23 | 0.154 |
|  | IV | 0.508 | 0.26 | 1.953 | 24 | 0.063 |
|  | V | 0.328 | 0.172 | 1.902 | 24 | 0.069 |
|  | VI | 0.183 | 0.133 | 1.378 | 24 | 0.181 |
| Treated vs Viral-Control : Post-Laser | **AVREC** | 0.402 | 0.196 | 2.054 | 24 | 0.051 |
|  | II | 0.478 | 0.423 | 1.961 | 23 | 0.062 |
|  | IV | 0.558 | 0.286 | 1.947 | 24 | 0.063 |
|  | V | 0.425 | 0.190 | 2.239 | 24 | **0.035** |
|  | VI | 0.232 | 0.147 | 1.581 | 24 | 0.127 |
| Treated : Post-Laser vs Pre-Laser | **AVREC** | 0.13 | 0.025 | 5.121 | 702 | **<0.001** |
|  | II | -0.052 | 0.046 | -1.132 | 729 | 0.258 |
|  | IV | 0.281 | 0.044 | 6.362 | 758 | **<0.001** |
|  | V | 0.115 | 0.038 | -4.597 | 754 | **0.003** |
|  | VI | 0.086 | 0.034 | 2.567 | 777 | **0.044** |

The control groups shared a similar pattern of activity in click train measurements, an increase rather than decrease in activity in the AVREC and thalamic input layers (Figure 3A). And significance was found in the CRY2olig and naïve control AVREC comparison and in the CRY2olig and viral control Layer V comparison (Table 1). Further post-laser group comparisons in the LMM were not found to be significant but did come close, for example at p values of 0.051 and 0.063 (Table 1). This is in contrast to the comparisons in amplitude modulation measurements, in which the control groups did not behave as comparatively and post-laser group comparisons with the CRY2olig group were far from significant (Figure 3B, Table 2).

**Table 2.** LMM results for Amplitude modulated stimulus: Comparisons run with the logRMS and the Intercept: CRY2olig-Treated:Post-Laser. Therefore the intercept, treated vs Naïve control post-laser, treated vs viral control post-laser, and treated pre-laser vs post laser are compared in the AVREC and layer traces. Significant results, p<0.05, are in bold.

| Predictors | Layer | Estimates | SE | Statistics | df | p |
| --- | --- | --- | --- | --- | --- | --- |
| Treated vs Naïve-Control : Post-Laser | **AVREC** | 0.187 | 0.169 | 1.106 | 24 | 0.28 |
|  | II | 0.188 | 0.195 | 0.964 | 23 | 0.35 |
|  | IV | 0.322 | 0.239 | 1.349 | 24 | 0.19 |
|  | V | 0.127 | 0.187 | 0.681 | 24 | 0.50 |
|  | VI | 0.113 | 0.161 | 0.7 | 24 | 0.49 |
| Treated vs Viral-Control : Post-Laser | **AVREC** | 0.258 | 0.187 | 1.382 | 24 | 0.18 |
|  | II | 0.543 | 0.214 | 2.924 | 23 | **0.02** |
|  | IV | 0.331 | 0.263 | 1.259 | 24 | 0.22 |
|  | V | 0.211 | 0.206 | 1.787 | 24 | 0.32 |
|  | VI | 0.205 | 0.177 | 1.155 | 24 | 0.26 |
| Treated : Post-Laser vs Pre-Laser | **AVREC** | 0.08 | 0.025 | 3.632 | 657 | **<0.001** |
|  | II | 0.134 | 0.046 | 2.924 | 693 | **0.004** |
|  | IV | 0.138 | 0.044 | 3.147 | 743 | **0.002** |
|  | V | 0.067 | 0.038 | 1.179 | 747 | 0.07 |
|  | VI | 0.077 | 0.033 | 2.315 | 775 | **0.02** |

The single control to CRY2olig group comparison which was found to be significant was the viral control vs the CRY2olig group in layer II. In the cortical response to click train measurements, the CRY2olig group shows a slight increase, non-significant after the laser which corresponds in directionality with the controls. In response to the amplitude modulation measurement, there is still a significant decrease in activity.

Despite the lack of significance between the CRY2olig and control groups in amplitude modulation measurements, it can still be observed that the CRY2olig group had a consistent decrease in activity after the laser which was found to be significant across both measurement types.

#### Single-trial t tests show that the effects between groups are much more significant when cortical recruitment is broad and intense

We ran single-trial Student’s *t* tests to follow the more conservative LMM in order to locate specific differences between all groups pre- and post-laser. Cohen’s d was calculated along with each *p* value to substantiate results. To begin with, within-group differences were found in control groups pre- and post-laser in click train measurements (Table 4). This implies that the heat from the laser could be amplifying the already ketamine-induced increase to recurrent excitation. A lack of differences found within control pre- and post-laser comparisons in amplitude modulation measurements attest to laser heat not being excessive and not affecting the less strongly recruited cortical responses. Significant differences were also found within the CRY2olig group pre- to post-laser which follows the LMM results (Figure 3).

**Table 3.**
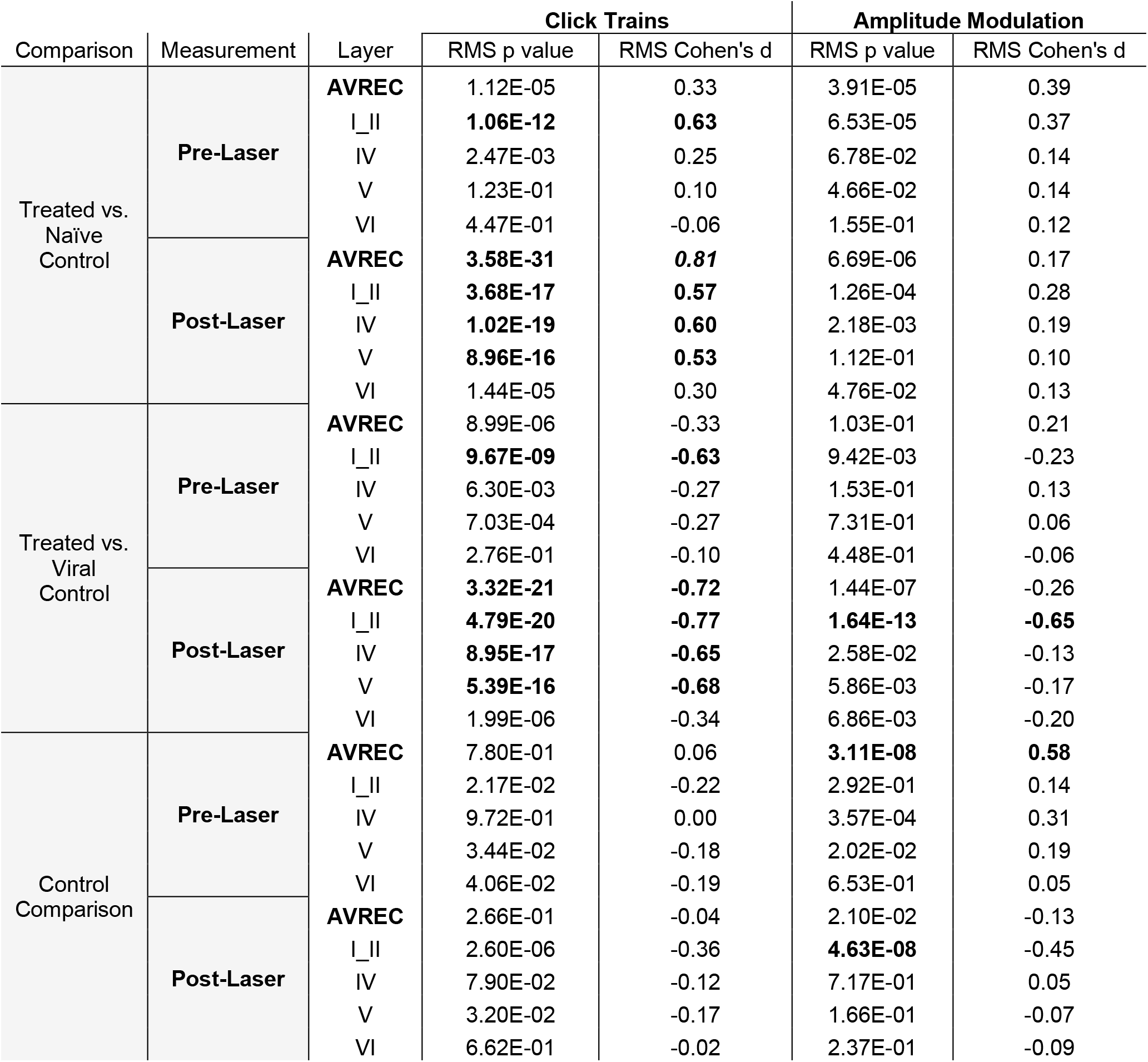
Between group comparisons: Treated vs Naïve control, Treated vs. Viral control, and Naïve control vs Viral Control comparisons during measurements taken pre- and post-laser in the full AVREC and layer traces (top to bottom). *P* and Cohen’s d results are shown for click train and amplitude modulated stimulations root mean square (RMS) for the first 200 ms. In bold are significance p < 1.00E-7 (Bonferroni corrected to 7.14E-08), corresponding to ❖ and ❖❖, as well as effect sizes over Medium d<0.5.

| Comparison | Measurement | Layer | Click Trains |  | Amplitude Modulation |  |
| --- | --- | --- | --- | --- | --- | --- |
|  |  |  | RMS p value | RMS Cohen's d | RMS p value | RMS Cohen's d |
| Treated vs. Naïve Control | Pre-Laser | AVREC | 1.12E-05 | 0.33 | 3.91E-05 | 0.39 |
|  |  | I_II | **1.06E-12** | **0.63** | 6.53E-05 | 0.37 |
|  |  | IV | 2.47E-03 | 0.25 | 6.78E-02 | 0.14 |
|  |  | V | 1.23E-01 | 0.10 | 4.66E-02 | 0.14 |
|  |  | VI | 4.47E-01 | -0.06 | 1.55E-01 | 0.12 |
|  | Post-Laser | AVREC | **3.58E-31** | **0.81** | 6.69E-06 | 0.17 |
|  |  | I_II | **3.68E-17** | **0.57** | 1.26E-04 | 0.28 |
|  |  | IV | **1.02E-19** | **0.60** | 2.18E-03 | 0.19 |
|  |  | V | **8.96E-16** | **0.53** | 1.12E-01 | 0.10 |
|  |  | VI | 1.44E-05 | 0.30 | 4.76E-02 | 0.13 |
| Treated vs. Viral Control | Pre-Laser | AVREC | 8.99E-06 | -0.33 | 1.03E-01 | 0.21 |
|  |  | I_II | **9.67E-09** | **-0.63** | 9.42E-03 | -0.23 |
|  |  | IV | 6.30E-03 | -0.27 | 1.53E-01 | 0.13 |
|  |  | V | 7.03E-04 | -0.27 | 7.31E-01 | 0.06 |
|  |  | VI | 2.76E-01 | -0.10 | 4.48E-01 | -0.06 |
|  | Post-Laser | AVREC | **3.32E-21** | **-0.72** | 1.44E-07 | -0.26 |
|  |  | I_II | **4.79E-20** | **-0.77** | **1.64E-13** | **-0.65** |
|  |  | IV | **8.95E-17** | **-0.65** | 2.58E-02 | -0.13 |
|  |  | V | **5.39E-16** | **-0.68** | 5.86E-03 | -0.17 |
|  |  | VI | 1.99E-06 | -0.34 | 6.86E-03 | -0.20 |
| Control Comparison | Pre-Laser | AVREC | 7.80E-01 | 0.06 | **3.11E-08** | **0.58** |
|  |  | I_II | 2.17E-02 | -0.22 | 2.92E-01 | 0.14 |
|  |  | IV | 9.72E-01 | 0.00 | 3.57E-04 | 0.31 |
|  |  | V | 3.44E-02 | -0.18 | 2.02E-02 | 0.19 |
|  |  | VI | 4.06E-02 | -0.19 | 6.53E-01 | 0.05 |
|  | Post-Laser | AVREC | 2.66E-01 | -0.04 | 2.10E-02 | -0.13 |
|  |  | I_II | 2.60E-06 | -0.36 | **4.63E-08** | -0.45 |
|  |  | IV | 7.90E-02 | -0.12 | 7.17E-01 | 0.05 |
|  |  | V | 3.20E-02 | -0.17 | 1.66E-01 | -0.07 |
|  |  | VI | 6.62E-01 | -0.02 | 2.37E-01 | -0.09 |

**Table 4.** Within group comparisons: Pre-vs post-laser comparison for Treated, Naïve control, and Viral Control groups in the full AVREC and layer traces (top to bottom). *P* and Cohen’s d results are shown for click train and amplitude modulated stimulations root mean square (RMS) for the first 200 ms.

| Group | Layer | Click Trains |  | Amp Mod |  |
| --- | --- | --- | --- | --- | --- |
|  |  | RMS p value | RMS Cohen's d | RMS p value | RMS Cohen's d |
| Treated | **AVREC** | 5.58E-04 | 0.256005 | 0.0187001 | 0.177 |
|  | I_II | 0.2656084 | -0.08769 | 0.0014983 | 0.248 |
|  | IV | 0.0003956 | 0.266303 | 0.1094612 | 0.121 |
|  | V | 0.0485516 | 0.146582 | 0.3142819 | 0.076 |
|  | VI | 1.62E-01 | 0.103797 | 0.2268568 | -0.09 |
| Naïve Control | **AVREC** | 0.1359423 | -0.10987 | 0.2974203 | 0.077 |
|  | I_II | 2.15E-01 | 0.092548 | 0.8882649 | 0.011 |
|  | IV | 0.6296055 | -0.03562 | 0.424591 | 0.06 |
|  | V | 0.000284 | -0.27004 | 0.3772673 | 0.066 |
|  | VI | 0.0213145 | -0.17007 | 0.4409822 | -0.06 |
| Viral Control | **AVREC** | 0.0274118 | -0.1944 | 4.234E-07 | -0.45 |
|  | I_II | 0.5258021 | -0.05651 | 0.0003823 | -0.32 |
|  | IV | 0.2966003 | -0.09275 | 0.0005953 | -0.31 |
|  | V | 2.12E-03 | -0.27317 | 0.0155829 | -0.22 |
|  | VI | 0.5901383 | 0.047402 | 0.0589803 | -0.17 |

Significance was found in click train measurements between CRY2olig and control groups pre-laser and post-laser in the AVREC, and all layers except pre-laser layer VI (Figure 4A). The magnitude of the *p* value difference is dramatically increased in post-laser comparisons (Table 3) and the Cohen’s d effect size increases from pre- to post-laser across the group comparisons as well, except in layer II. Cohen’s d is described in more detail below.

**Figure 4.**
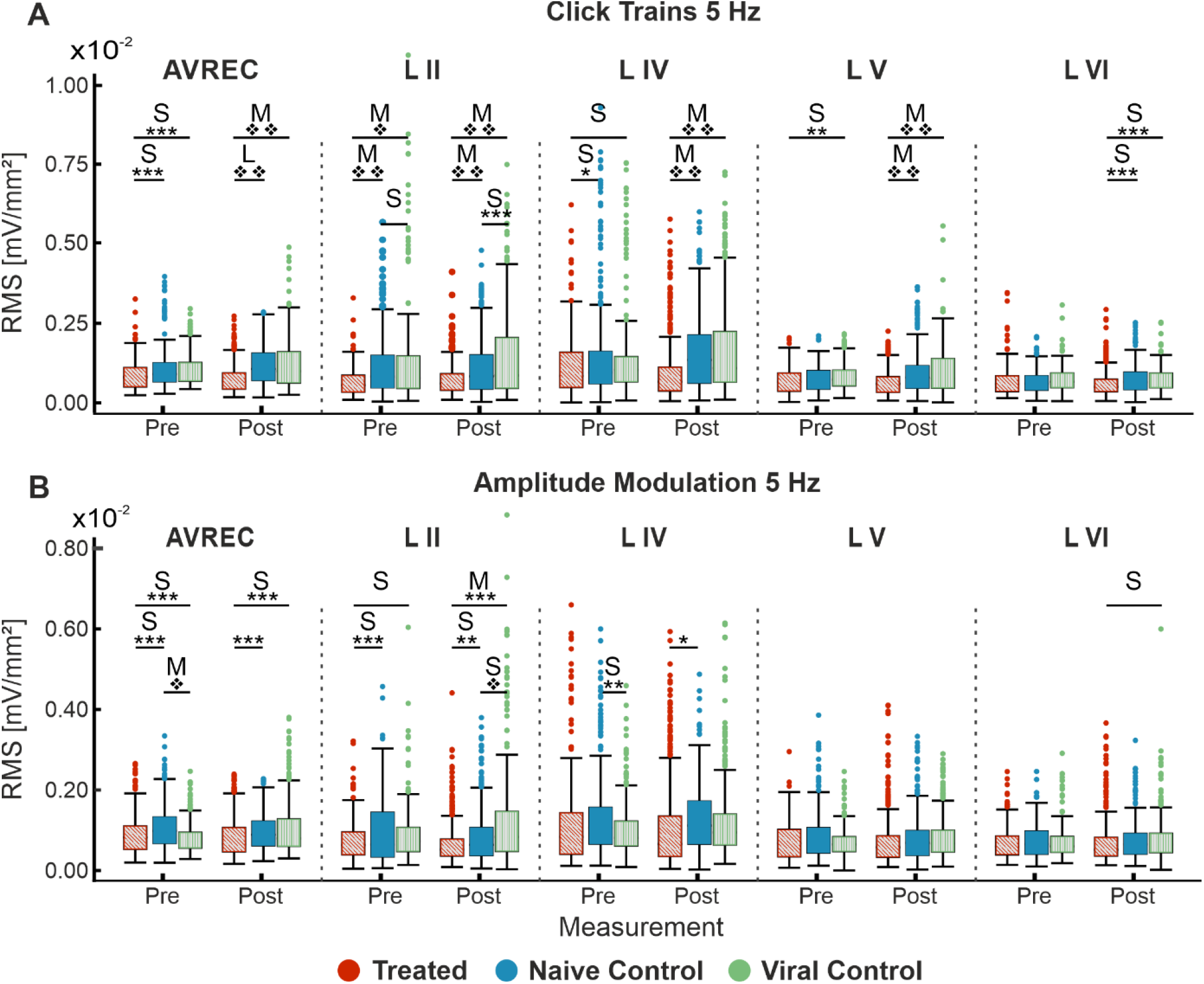
Single-trial box plots for 5 Hz; **A**. Single-trial box plots for CRY2olig-treated, naïve, and viral control groups for the AVREC and thalamic input layers II, IV, V, and IV RMS in response to the first 200 ms of 5 Hz click trains. B. Single-trial box plots for CRY2olig, naïve, and viral control groups for the AVREC and thalamic input layers II, IV, V, and IV RMS in response to the first 200 ms of 5 Hz amplitude modulation. Student’s *t* test and Cohen’s d effect size results overlaid when significant or at least small, respectively. p < 0.05 *, <0.01 **, <0.001 ***, <0.000001❖, <1E-10❖❖, Bonferroni corrected in single-trial comparisons (n=14). Cohen’s d 0.2-0.5 = small, 0.5-0.8 = medium, 0.8-1.2 = large.

For amplitude modulation measurements, there was significance found pre- and post-laser between the CRY2olig and at least one of the control groups for the AVREC and layer II (Figure 4B, Table 3). Only small effect sizes were found in the comparison with between the CRY2olig and control groups in the AVREC or thalamic layers. There was a medium effect size difference and significance found before the laser between the control groups in the AVREC.

The viral control group appeared the most abnormal compared to cortical activity in the naïve control group for amplitude modulation. However, the directionality of change pre- to post-laser for the viral control group between click train and amplitude modulation was consistent (increase after laser) while the naïve control group had a slight decrease after laser in amplitude modulation measurements in contrast with its increase after laser in the click train measurements. Again, the CRY2olig group cortical activity consistently decreased after the laser, although the magnitude of significance was much less in the amplitude modulation pre- and post-laser comparisons.

#### Cohen’s d effect sizes clarify magnitude changes and highlight clustering effects differentiating the CRY2olig from control groups during click train stimuli

Cohen’s d effect size was calculated along with each *t* test. The Cohen’s d effect sizes quantitatively motivate a more wholistic understanding of this data and its implications. A stark increase was found in AVREC effect sizes pre- and post-laser for cortical response strength to 5 Hz click trains (Figure 5A) in both CRY2olig vs control comparisons. And there was a 1-fold or 2-fold increase in thalamic input layers in effect size of difference between CRY2olig and control groups after the laser presentation.

**Figure 5.**
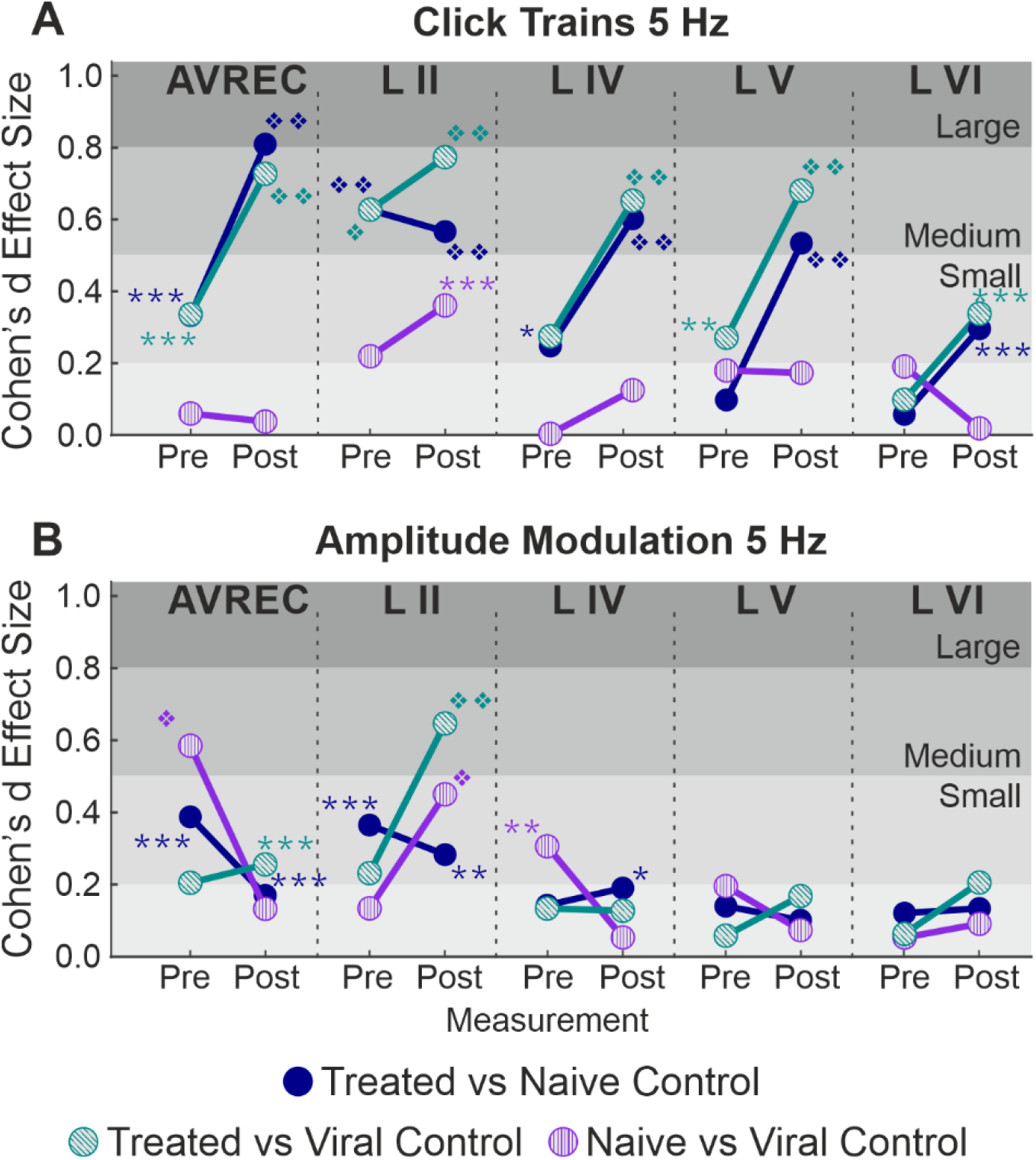
Cohen’s d effect sizes; All single-trial tests included. **A**. 5 Hz click train Cohen’s d effect sizes and overlaid student’s *t* test p value results for comparison between CRY2olig and naïve control (blue), CRY2olig and viral control (green), and the naïve and viral control groups (purple) both before and after the laser. Values compared were the RMS of the first region of interest time window of response for the AVREC trace and all layer traces (left to right). **B**. 5 Hz amplitude modulation Cohen’s d effect sizes and overlaid student’s *t* test p value results for comparisons between the treat and control groups before and after the laser for the AVREC and layer traces. p < 0.05 *, <0.01 **, <0.001 ***, <0.000001❖, <1E-10❖❖, Bonferroni corrected. Cohen’s d 0-0.2 = negligible, 0.2-0.5 = small, 0.5-0.8 = medium, 0.8-1.2 = large.

CRY2olig vs. control group comparisons show medium or large effect sizes post-laser in all layers except layer VI while the pre-laser comparisons are small or negligible except in the case of the supragranular layer which was the most variable area between and within animals. Control comparisons across click train responses remain small and negligible in effect size.

The cortical response to amplitude modulation at 5 Hz (Figure 5B) showed small and negligeable effect sizes in all cases except the supragranular layer and the AVREC.

Most notable, is the directionality of effect size change between stimulus types. CRY2olig to control click train comparisons—in all but layer II—show an overwhelming increase in effect size and, in many cases, an increase in the order of magnitude of *p* value results after laser presentation. This couples with the consistent decrease in activity after the laser within the CRY2olig group and a general increase in activity after the laser in control groups. However, the amplitude modulation results indicate most often a decrease in Cohen’s d effect size after laser presentation which argues against the validity of post-laser *p* value results found. These *p* values are also of much lower magnitude in all cases except in the supragranular layer II and in this, the viral control group is the outlier with CRY2olig vs viral control and naïve vs viral control post-laser comparisons having higher effect sizes and very high magnitude *p* values.

### Spontaneous activity indicated systemic changes due to the presence of CRY2olig

Spontaneous activity was recorded while no stimuli were presented to explore the hypothesis that systemic effects of VGCC aggregation would be ongoing. Figure 6A and B show that there is no suppression of activity from pre to post laser in the treated group (except in Layer V). However, there was a highly significant group difference of RMS over the 1400 ms bins (of over 2 minutes of spontaneous activity per session), in single trial analysis, largely independent of laser stimulation (Figure 6B and C, LMM results in Table 5, between-group comparisons in Table 6). With a strong significant difference in the AVREC and layer traces between the CRY2olig and control groups, the only scale increase in significance pre- to post-laser was the CRY2olig and naïve control group in layer VI. Effect sizes were Large or Medium in most CRY2olig vs control comparisons, especially in the overall AVREC and in layer II. Control comparisons were also significantly different, with at most a Small effect size, in most cases but, notably, layer II and VI result in a negligeable effect size and no significant difference pre- or post-laser. Within-group comparisons of pre- to post-laser spontaneous activity (Table 7) found only one result of mild significance—in the naïve control group pre-laser—and negligeable effect size across the board. The LMM (Figure 6B) also confirmed that the CRY2olig group had a lower level of activity already before the laser in the AVREC, and layer traces II, IV, and VI, did not further suppress after laser presentation.

**Table 5.** results for Spontaneous Activity: Comparisons run with the logRMS and the Intercept: CRY2olig-CRY2olig:Post-Laser. Therefore the intercept, CRY2olig vs Naïve control post-laser, CRY2olig vs viral control post-laser, and CRY2olig pre-laser vs post laser are compared in the AVREC and layer traces. Significant results, p<0.05, are in bold.

| LMM Results: Spontaneous Activity |  |  |  |  |  |  |
| --- | --- | --- | --- | --- | --- | --- |
| Predictors | Layer | Estimates | SE | Statistics | df | p |
| CRY2olig vs Naïve-Control : Post-Laser | **AVREC** | 0.312 | 0.230 | 1.358 | 19 | 0.190 |
|  | II | 0.388 | 0.254 | 1.527 | 19 | 0.143 |
|  | IV | 0.415 | 0.343 | 1.209 | 19 | 0.242 |
|  | V | 0.164 | 0.297 | 0.552 | 19 | 0.587 |
|  | VI | 0.208 | 0.179 | 1.159 | 19 | 0.261 |
| CRY2olig vs Viral-Control : Post-Laser | **AVREC** | 0.431 | 0.242 | 1.779 | 19 | 0.091 |
|  | II | 0.336 | 0.268 | 1.253 | 19 | 0.225 |
|  | IV | 0.625 | 0.363 | 1.723 | 19 | 0.101 |
|  | V | 0.277 | 0.313 | 0.884 | 19 | 0.388 |
|  | VI | 0.235 | 0.190 | 1.241 | 19 | 0.230 |
| CRY2olig : Post-Laser vs Pre-Laser | **AVREC** | 0.013 | 0.012 | -1.085 | 1522 | 0.278 |
|  | II | -0.022 | 0.014 | -1.538 | 1522 | 0.124 |
|  | IV | 0.014 | 0.019 | 0.722 | 1522 | 0.470 |
|  | V | -0.044 | 0.016 | -2.666 | 1522 | **0.008** |
|  | VI | -0.00 | 0.013 | -0.015 | 1522 | 0.988 |

**Table 6.** Between group spontaneous AVREC and layer trace comparisons: CRY2olig vs Naïve control, CRY2olig vs. Viral control, and Naïve control vs Viral Control comparisons during measurements taken pre- and post-laser in the full AVREC and layer traces (top to bottom). *P* and Cohen’s d results are shown for spontaneous activity root mean square (RMS) for 1400 ms bins. In bold are significance p < 1.00E-7 (Bonferroni corrected to 7.14E-08), corresponding to ❖ and ❖❖, as well as effect sizes over Medium d<0.5.

| Comparison | Measurement | Layer | Spontaneous Activity |  |
| --- | --- | --- | --- | --- |
|  |  |  | RMS p value | RMS Cohen's d |
| CRY2olig vs. Naïve Control | Pre-Laser | AVREC | **2.12E-22** | **0.63** |
|  |  | I_II | **3.00E-39** | **0.77** |
|  |  | IV | 0.00012 | 0.27 |
|  |  | V | 0.066 | 0.12 |
|  |  | VI | **1.18E-10** | 0.41 |
|  | Post-Laser | AVREC | **1.81E-31** | **0.79** |
|  |  | I_II | **9.60E-45** | **0.80** |
|  |  | IV | 0.00015 | 0.28 |
|  |  | V | 0.00034 | 0.24 |
|  |  | VI | **1.41E-21** | **0.62** |
| CRY2olig vs. Viral Control | Pre-Laser | AVREC | **1.20E-34** | **-0.81** |
|  |  | I_II | **1.04E-24** | **-0.66** |
|  |  | IV | **2.78E-16** | **-0.53** |
|  |  | V | 9.52E-08 | -0.35 |
|  |  | VI | **7.13E-14** | **-0.50** |
|  | Post-Laser | AVREC | **2.26E-39** | **-0.88** |
|  |  | I_II | **3.70E-29** | **-0.73** |
|  |  | IV | **6.34E-18** | **-0.57** |
|  |  | V | **5.29E-11** | -0.43 |
|  |  | VI | **1.43E-22** | **-0.67** |
| Control Comparison | Pre-Laser | AVREC | **6.53E-11** | -0.42 |
|  |  | I_II | 0.173 | -0.08 |
|  |  | IV | **1.83E-11** | -0.45 |
|  |  | V | 5.16E-06 | -0.29 |
|  |  | VI | 0.074 | -0.11 |
|  | Post-Laser | AVREC | **9.81E-09** | -0.37 |
|  |  | I_II | 0.195 | 0.08 |
|  |  | IV | **9.18E-14** | -0.499 |
|  |  | V | 8.65E-06 | -0.28 |
|  |  | VI | 0.312 | -0.06 |

**Table 7.**
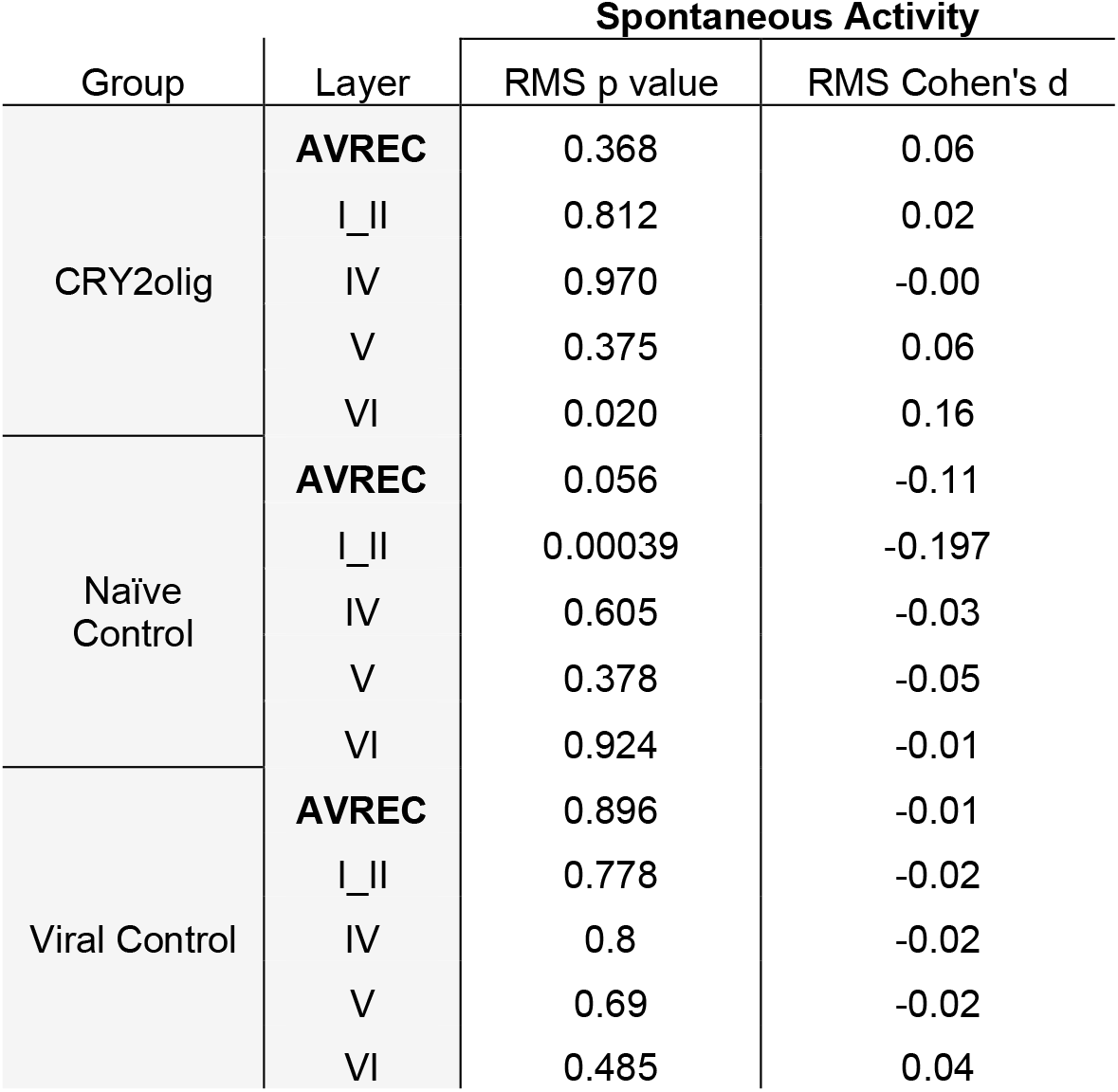
Within group AVREC and layer trace comparisons: Pre- vs post-laser comparison for CRY2olig, Naïve control, and Viral Control groups in the full AVREC and layer traces (top to bottom). *P* and Cohen’s d results are shown for spontaneous activity root mean square (RMS) for 1400 ms bins.

**Figure 6.**
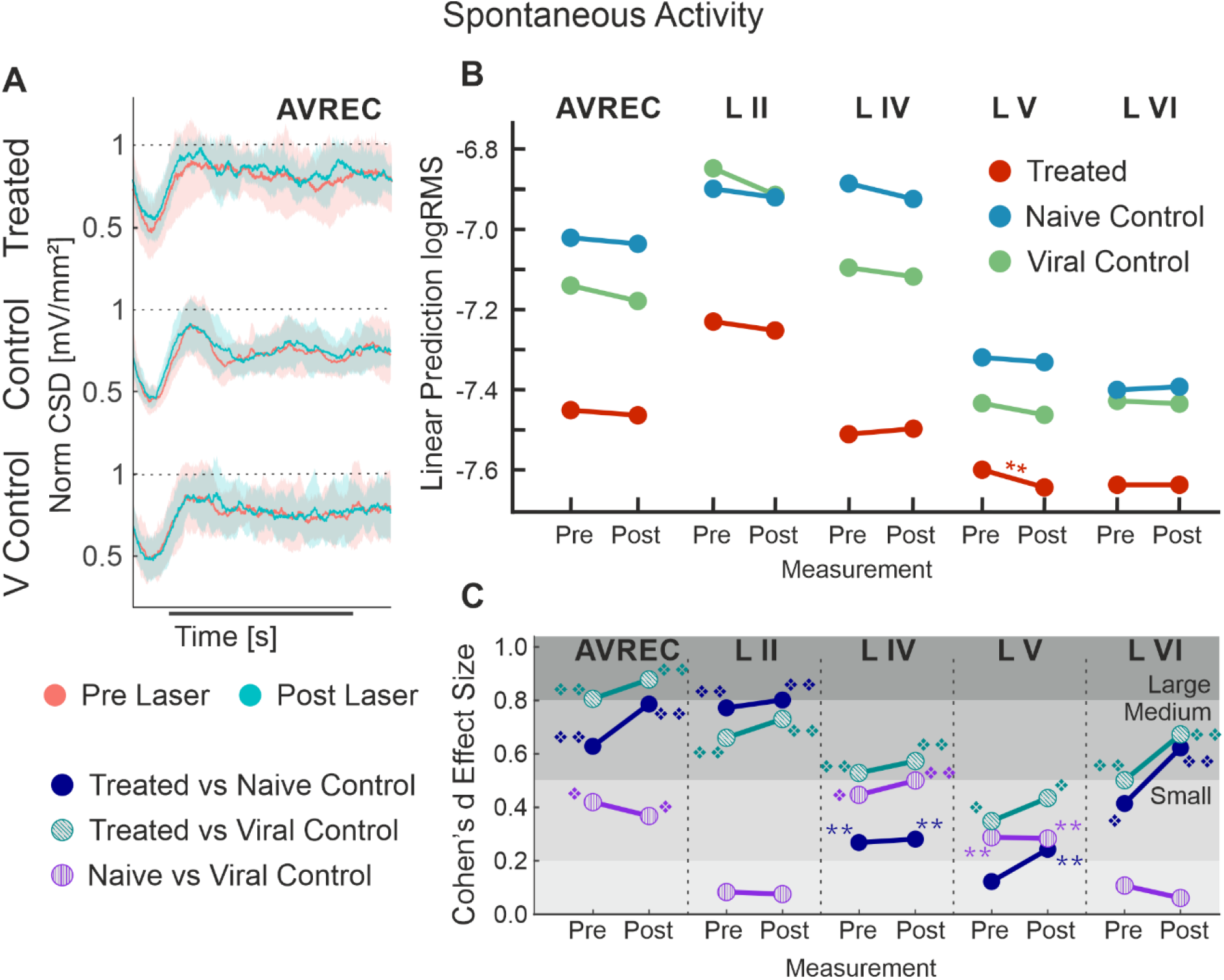
AVREC spontaneous activity; **A**. Grand averaged (±STD) AVREC traces of Cry2olig-treated (top), naïve control (middle), and viral control (bottom) subjects without stimulus presentation before (peach) and after (cyan) laser presentation. **B**. LMM effect plots for linear prediction over measurement (pre- to post-laser) for CRY2olig-treated (orange), naïve control (blue), and viral control (green) groups across the AVREC and all cortical layers without stimulus presentation. **C**. Cohen’s d effect size plot for the AVREC and layer comparison CRY2olig vs naïve control (dark blue), CRY2olig vs viral control (teal), and naïve vs viral control (purple). p < 0.05 *, <0.01 **, <0.001 ***, <0.000001❖, <1E-10❖❖, Bonferroni corrected for single trial comparisons. Cohen’s d 0-0.2 = negligible, 0.2-0.5 = small, 0.5-0.8 = medium, 0.8-1.2 = large.

### Temporal precision and synchronicity under ketamine anesthesia

In order to further understand the relationship that VGCC clustering across a population might have with internal synchronicity dynamics, vector strength analysis and spectral analysis were both calculated. Vector strength analysis was performed to understand the relationship of temporal precision of population response to amplitude modulation and click train stimuli. Both 5 Hz click trains and amplitude modulation encourage the cortical response to be highly synchronous (∼0.8 vector strength) while cortical response is asynchronous with both 10 Hz click trains and amplitude modulation (∼0.3 vector strength). The level of temporal precision does not change before and after laser presentation or over the next hour after laser presentation (ANOVA measurement differences: ns). There are some slight group differences in the activity in response to 5 Hz click trains and 10 Hz amplitude modulation, but there are no interaction effects (data not shown).

Spectral analysis was also run and both power and phase coherence were compared by permutation testing after continuous wavelet analysis and the resultant time frequency plots (Deane et al. 2020; Deliano et al. 2020). There was no difference found between groups in each condition (data not shown).

Analysis of spontaneous activity, recordings of cortical activity under anesthesia when no stimuli were present, also did not yield differences pre- to post-laser or between groups, suggesting that internal dynamics of ongoing synchronization were not different under ketamine anesthesia (data not shown).

## Discussion

The firing variability of neuronal populations, and thereby their evoked activity levels, partly depends on the lateral mobility of CaV2.1 voltage-gated calcium channels (VGCCs). In cultured primary hippocampal neurons, VGCC clustering with Cry2olig led to increased initial responses to electrical pulses, and a significant paired-pulse depression (Heck et al. 2019; Taslimi et al. 2014). In this study, we have combined the usage of CRY2olig with a transgenic mouse model (Mark et al. 2011) that allows for the optogenetic clustering of VGCCs *in vivo* to investigate the impact of VGCC aggregation in the primary auditory cortex (A1). Using different auditory stimulus classes: click trains and AM tones, we recorded laminar current source density (CSD) profiles in order to investigate the synaptic population activity within distinct cortical layers.

We found that VGCC clustering in the A1 led to a significant reduction in activity across both stimulus types. Clustering during the click train stimulus in particular saw significantly different activity levels between CRY2olig and control groups. Under anesthesia, cortical responses to sensory stimuli are generally enhanced and more reliable compared to the awake state (Deane et al. 2020). The aggregation of VGCC’s reduced the ability of cortical circuits to amplify input information. Our findings thereby provide evidence that synaptic variability instigated by presynaptic VGCC membrane motility serves as an important maintenance in the encoding of sensory signals by dynamically adjusting network activity.

### VGCC clustering time course and laser-heat have opposing effects

The overall activity level in the CRY2olig group was already suppressed compared to the controls before the laser, especially in click train measurements (Figure 3) and spontaneous activity (Figure 6). Additional and significant activity suppression after the laser light was observed for over an hour following the VGCC clustering in response to stimulus presentation. This effect extended beyond the time-course for channel clustering measured *in vitro*, which is ∼60 % directly after light stimulation and decreases to ∼30 % over 30-40 minutes and to ∼0 % again in the duration of 160 minutes (see Taslimi et al., 2014 Fig. 1, Heck et al. 2019). Therefore, we conclude that this pre-laser and spontaneous suppression compared to controls is potentially due to a light-activated clustering during the surgical and experimental procedures. Contrastingly in the control groups, we found increased activity after the laser illumination. Laser-induced increases in neuronal activity have been observed in multiple studies (Arias-Gil et al. 2016; Brunk et al. 2019; Owen et al. 2019; Stujenske et al. 2015). The systematic suppression effect of the VGCC clustering consistently counteracted this laser-induced increase. Therefore, our methodological approach, if anything, underestimates the effect strength of VGCC clustering on sound-evoked cortical responses.

The level of cortical activity during spontaneous recordings remained constant throughout experiments, with very little dependance on the laser. This indicated that VGCC clustering resulted in a systemic, long-term change. In the click train and amplitude modulated responses, there was clear suppression after laser stimulation. Stimulus response circuits using the affected networks where seemingly suppressed to different levels depending on the amount of circuitry used. As in, in response to the highly synchronized, tonotopically activating, click trains, thalamic input layers IV and V were more strongly suppressed after the laser than in response to the narrow spectrum, amplitude modulated pure tone. Therefore, refreshing VGCC aggregation with laser presentation further suppressed responses to amplitude modulated pure tones and click trains in a level and laminar dependent way.

### Columnar suppression of impulse responses due to VGCC clustering

We compared the CRY2olig group pre- to post-laser and between groups post-laser with a linear mixed model (LMM, Figure 3). Results indicated that pre- to post-laser suppression of activity was significant across most layers after VGCC clustering across both stimulus types. Contrasting the VGCC-induced suppression, there was an overall tendency in the control groups toward increased activity after laser stimulation. The LMM analysis revealed significant between-group comparisons after the laser for some of the most pronounced effects on cortical layer activity. Specifically, VGCC clustering caused a significant reduction of the overall columnar response strength measured by the AVREC and sound-evoked synaptic activity in cortical layer V during the click train stimulation, and significantly lower activity in layer II during AM-stimulation. VGCC aggregation had differential effects on sensory processing of stimulus classes that cause broad spectral and highly synchronized thalamocortical synaptic input, compared to population activity, which relies more on temporal integration of intracortical synaptic inputs.

To further tease apart group differences, single-trial, Bonferroni corrected, Student’s *t* tests were calculated pre- and post-laser between all groups (Figure 4). Results were contextualized with Cohen’s *d* effect sizes (Figure 5). Responses to click trains were already significantly different between CRY2olig and control groups before laser-induced VGCC aggregation for the AVREC, and sound-evoked activity recorded in cortical layers IV and V. Effects increased in magnitude of significance as well as in effect size after laser presentation. Amplitude modulation comparisons yield much lower significance, when found, post-laser, and only had a medium effect size in post-laser layer II and in the pre-laser AVREC control comparison. The less strong synchronized recruitment of synaptic populations with this stimulus class most likely explains the less prominent effects.

Our results indicate that the clustering presynaptic VGCC is detrimental to overall population activity— hence the suppression—which is exacerbated in circumstances of high cortical recruitment. Such recruitment of recurrent excitation circuits is found particularly during the representation of salient and behaviorally relevant stimuli (Kato et al. 2017). Additionally, recurrent excitation in thalamic input layers of sensory cortex may play a central role especially for enhancing threshold-near stimuli (Happel and Ohl 2017; Wang 2008). Our study now hints that the neural basis of such circuit-derived enhancement in the sensory cortex may be at least partly rooted in the presynaptic dynamics of VGCCs.

### Amplification disruption of thalamic input layers

Recurrent excitation in layer IV has been implicated as the dominant circuit activity contributing to the AVREC response (Deane et al. 2020; Happel et al. 2010). It is henceforth unsurprising that this and thalamic input layer V most closely resembled the AVREC in the click train measurement cortical responses (Figure 3A). In single-trial group comparisons, there was a two-fold increase in effect size and an overwhelming increase in the magnitude of significance in the AVREC pre- to post-laser. This two-fold increase is reflected in layer V and a one-fold increase is found in layers IV and V.

The reduction of impulse responses and ongoing responses after click train stimulation (Figure 2) can therefore be explained by the fact that the stochastic firing variability of individual synapses, reduced by the clustering of VGCCs, plays an important role in recurrent excitation in layer IV. By aggregating VGCCs we change the temporal resolution of recurrent excitation and, therefore, disrupt the gain function of cortical amplification circuits.

### Dynamic supra- and infragranular responses

Click-train evoked responses in supragranular layers showed an opposing trend compared to the other layers: VGCC clustering led to a slight increase of activity. This is consistent with AM stimulation (Figure 3B) where we observed the decrease in supragranular activity after the laser in the CRY2olig group, consistent with the overall suppression after VGCC clustering. Pre-laser activity in the controls is generally lower in this layer during AM tones compared to click trains, owing likely to a high volume of cross-columnar activity through the dense network in supragranular layers following click stimuli. While the heat from the laser, described above, would be most intense on the surface of the cortex, it did not cause an increase in activity across both types of stimulation in supragranular layers. This suggests that the effect of the laser, the VGCC clustering, and the higher recruitment of click trains coincide in a non-linear fashion. What might be concluded from this is that the network is more sensitive to light effects during high recruitment than clustering effects, therefore causing the increase in activity after clustering in the click train condition, but that suppression after clustering is more broadly consistent across different conditions (Figure 3).

Layer VI in the cortical microcircuit is largely seen as the main feedback to the thalamus, completing a cortico-thalamic loop that has been discussed with respect to cortical gain during sensory processing and perception (Alitto and Usrey 2003; Homma et al. 2017; Saldeitis et al. 2021). While these layers are generally less active under anesthesia, during click train cortical responses, there was a significant increase in cortical activity after VGCC clustering (Figure 3A, LVI). Effects on aggregation in other layers may have caused a disinhibition of synaptic activity in deeper layers, explaining these opposing effects. Such disinhibitory feedback on the cortical gain has been related to Layer IV corticothalamic neurons in the auditory cortex (Williamson and Polley 2019).

## Conclusion

In this study, we have used CRY2olig to aggregate VGCCs in the A1 of transgenic mice. By using laminar CSD analysis *in vivo*, we could show that manipulation of lateral membrane motility of VGCCs in the presynaptic terminals significantly modulates population activity in all cortical layers. Our results indicate a more general loss of function in sensory processing due to the aggregation of these channels, despite an artificially created increase in initial firing response at a single-unit level (cf. Heck et al., 2019). In comparing results between click train stimuli, AM tones, and spontaneous activity, we argue that this loss of functionality is most critical in cases of strong cortical recruitment due to highly synchronized synaptic inputs—a key feature of recurrent excitation in sensory cortex during processing of salient, behaviorally relevant sensory cues.

## Additional Information

### Author contributions

Research was performed in the laboratories of the Leibniz-Institute for Neurobiology, Magdeburg (Germany), and designed by KED, JH, MH, and MFKH with input from FWO. MFKH supervised the project. The mouse line was contributed by MDM. Experiments were performed by KED. Data and statistical analysis were performed by KED and RK. Figures were prepared by KED. KED wrote the initial paper draft. Manuscript was written and edited by KED, RK, and MFKH with assistance from JH, MH, and FWO. All authors reviewed the manuscript, approve of the final version, and agree to be accountable for all aspects of the work. All persons designated as authors qualify and all who qualify are listed as authors.

## Acknowledgements

We would like to thank Kathrin Ohl and Anja Gürke for their technical assistance.

